# Effect of eco-remediation and microbial community using multilayer solar planted floating island (MS-PFI) in the drainage channel

**DOI:** 10.1101/327965

**Authors:** Zhen Sun, Dongge Xie, Xiran Jiang, Guihong Fu, Dongxue Xiao, Liang Zheng

## Abstract

A multilayer solar planted floating island (MS-PFI) planted with *Eichhornia* crassipes are potential alternatives to traditional PFI. The highest removal rates of suspended solids, total nitrogen, total phosphorus, ammonia nitrogen and chemical oxygen demand was 86%, 75%, 80%, 95% and 84%, respectively. Proteobacteria (average 43.4% of total sequences) and Actinobacteria (19.9%) were the dominant phyla. Numerous genus had obvious differences between influent and effluent water, for instance, 13, 12 and 7 % in effluent water were assigned to the hgcl_clade, Norank_c_Cyanobacteria, and *Rhizorhapis*, while their relative abundances were decreased to 5, 3 and 0 %. In contrast, a distinct increase among *Flavobacterium* (10%), *Limnohabitans* (7%), *Alpinimonas* (4%), norank_p_Saccharibacteria (4%), *Erwinia* (3%) after MS-PFI treatment. MS-PFI brings various bacteria involved in contaminant degradation and nutrient removal in biological wastewater treatment systems. An amount of ¥ 1,843 was totally inputted to construct floating bed, which was rarely needed operation and maintenance costs.

**Importance:** In-situ micro-polluted water ecological remediation, microorganisms and plants are effective to improve environmental quality and provide essential ecosystem services. Recently, we invent a new multilayer solar with an excellent pollutant removal efficiency. Microbes can decompose or mineralize organic matter effectively, also provide food for aquatic animals and increase nutrients or substances for plants, it is an important part of biogeochemical cycles and energy flows in aquatic ecological systems. However, few study explain the bacteria diversity and its responses between influent and effluent water in a planted floating island. The significance of our study is in identifying-in greater detail-the responses of bacteria in the new MS-PFI. This will greatly enhance our knowledge of bacteria communities, and can be widely used in micro-polluted water remediation.

## Introduction

With the rapid development of industrial production and population in the past few decades, a growing number of pollutants were discharged into the surface water without appropriate treatment, which can cause a huge threat to the quality of drinking water sources and the health of citizens. This contaminated surface water is referred to as micro-polluted surface water (MPSW) (1). Generally, it contains chemical oxygen demand COD, nitrogen, phosphorus and carbon pollutants are of relatively low concentrations (that is why it is called micro-pollution). Majority of indexes are no more than 10 mg/l, and this makes a specially designed for water treatment process (2). Since the 1980s, planted floating island has been widely researched and applied in USA (3,4), Netherlands (5), Japan (6), China (7, 8). Numerous scientists (9,10) summarized the design, operation, plant, and management of planted island treatment, especially the data meta-analysis of various applications. Remarkably, *Eichhornia* crassipes have been widely applied in a floating island for the treatment of surface water and wastewater, for the ability to adapt equally well at several location and climate zone (11, 12, 13, 14).

Plants and microorganisms are important ecological factors of MS-PFI, due to the efficacy in assimilating nutrients and in creating favorable conditions for the decomposition of organic matter. Assimilation is fixed in plants, allowing the harvest of plants to remove nitrogen and phosphorus from the aquatic environment. At the same time, the roots and foam carriers can attract largely microbes, increasing energy and oxygen to benefit the microbes, completing the degradation function (15). Microbes can decompose or mineralize organic matter effectively, also provide food for aquatic animals and increase nutrients or substances for plants (16). Microbe is an important part of biogeochemical cycles and energy flows in aquatic ecological systems. A complete analysis of bioprocess needs the characterization of the microbial communities developing in the constructed island, mainly those belonging to Bacteria domain (17). In this sense, it is urgent to explore bacterial communities thriving in influent and effluent water among water treatment systems. For its ability for deep sequencing of bacterial communities and identification of rare populations in low abundance, high-throughput sequencing technology has become a popular tool for examining bacteria communities (18).

This paper was developed a new MS-PFI, which combined the advantageous components for traditional PFI, in addition to a solar battery system to increase oxygen by lifting and circulating water. We specifically investigated: (1) the efficiency of water purification by MS-PFI; (2) the characteristics of the bacterial community in influent and effluent water samples of MS-PFI; (3) economic benefit evaluation of MS-PFI. Our design and experimental data provided the future applications of MS-PFI in lakes, reservoirs, and artificial landscape water to help solve the problems that caused by conventional devices, and without consuming extra energy sources.

## Results

### Investigation of reference area

The reference samples collected near a sewage draining exit to investigate the water quality in campus sewage background. The range and the average values of water quality shown in Table 1. The average water temperature was about 22.6°C, which was consistent with the environmental temperature from April to June. Many water quality indexes were very high, such as COD_Cr_ and TN. Campus ditches received a high load of domestic wastewater could cause biodiversity decrease and water quality deterioration. Thus, it is necessary to strengthen the wastewater treatment from campus water in this area.

**Table 1.** The quality indexes of the reference area

| Index | Range | Average |
| --- | --- | --- |
| Water Temperature (°C) | 19.0~25.2 | 22.6 |
| pH | 7.36~7.90 | 7.68 |
| SS (mg/L) | 1.0~3.0 | 2.4 |
| TN (mg/L) | 0.48~2.32 | 1.17 |
| TP (mg/L) | 0.01~0.04 | 0.03 |
| NH<sub>3</sub>-N (mg/L) | 0.00~ 0.04 | 0.01 |
| COD<sub>Cr</sub> (mg/L) | 23.808~ 59.520 | 35.712 |

### The effect of plant floating island

This study used grown *Eichhornia* crassipes on a floating island to test potential in phytoremediation water bodies. Table 2 summarized the significant statistical results for the t-test of paired differences, including mean, Std Deviation, Std Error mean, t statistic and p-value. All water quality indexes were significantly different between JW and CW for all p-value >0.05, which means the water quality had a great improvement after MS-PFI treatment. MS-PFI had apparent stratification in water temperature and pH prior to the first day in the experiment (Fig. 2). The average water temperature was 22.7±0.3°C in the MS-PFI effluent samples, 2.3 degrees lower (P=0.05) than the influent temperature (25.0±0.4°C). A small increase was observed in pH range from 7.66±0.01 to 7.77±0.01 (Fig. 2). pH existed difference ranges from 0.07 to 0.15, which became more stable about 0.11 after 24 days.

**Table 2.** Paired sample test between JW and CW treatment

| Pair items | Mean | Std.<br>Deviation | Std. Error<br>Mean | t | df | Sig<br>(2-tailed) |
| --- | --- | --- | --- | --- | --- | --- |
| JW-CW water temperature | 2.30700 | 1.23868 | .39170 | 5.890 | 9 | .000 |
| JW-CW pH | -.10867 | .02903 | .00918 | -11.839 | 9 | .000 |
| JW-CW SS | 2.78024 | .30565 | .09665 | 28.765 | 9 | .000 |
| JW-CW TP | .02337 | .01330 | .00421 | 5.556 | 9 | .000 |
| JW-CW TN | .72894 | .51803 | .16382 | 4.450 | 9 | .002 |
| JW-CW NH<sub>3</sub>-N | .03410 | .02789 | .00882 | 3.866 | 9 | .004 |
| JW-CW COD | 18.91413 | 10.90788 | 3.44937 | 5.483 | 9 | .000 |

**Fig 1.**
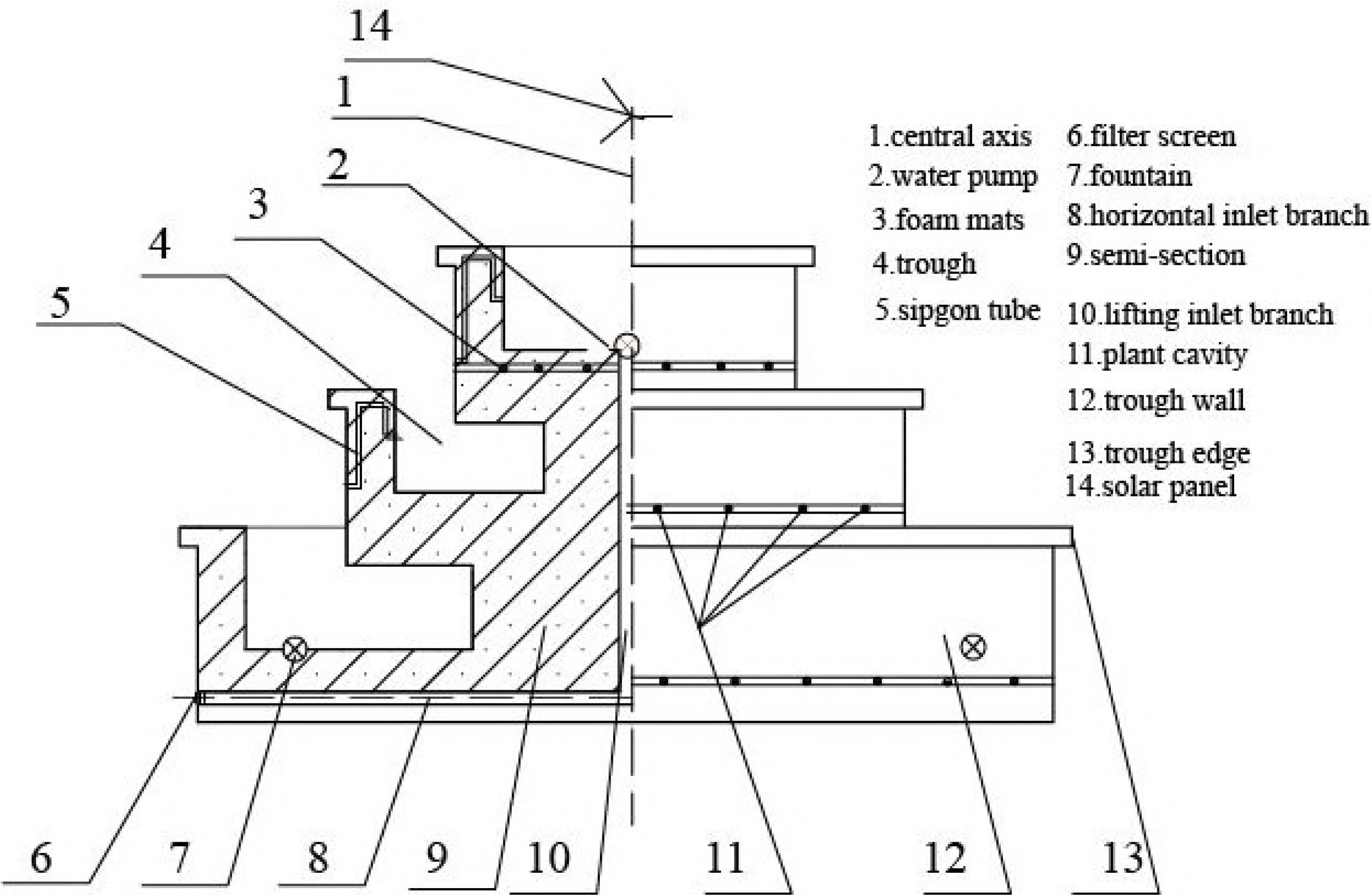
Design drawing of multilayer solar plant floating island water purification system

**Fig 2.**
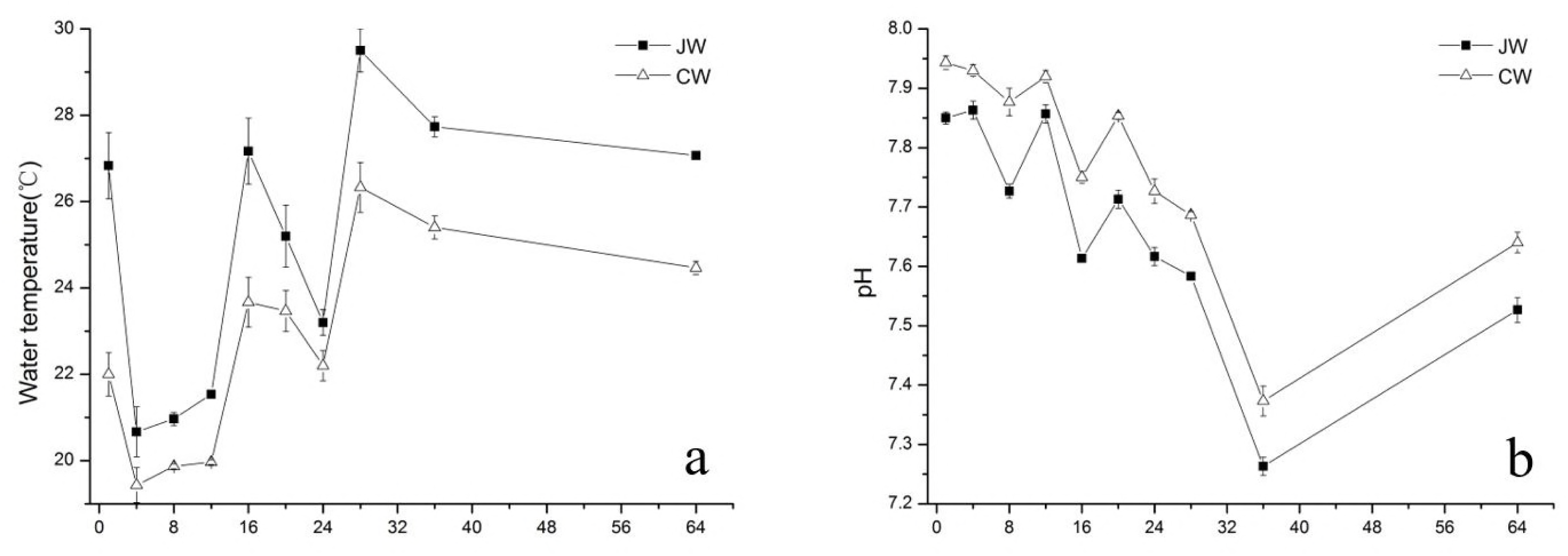
Water temperature and pH were detected in influent and effluent samples of MS-PFI

The removal rate of SS, TN, TP, NH_4_^+^-N, and COD were 67%, 50%, 51%, 69% and 61%, respectively (Fig. 3). The SS concentration in influent samples stayed high, while SS in effluent samples remained relatively low, namely, 3 and 0 mg/L respectively. The removal rate of SS was efficient on all days, the highest removal rate was 87%. However, the lower SS removal rate appeared in the early period of treatment (for example, the first week). The average concentration of TP was 0.05mg/L in influent water, then reduced by 51% in effluent water. TN concentration decreased by the effect of MS-PFI, the average removal rate of TN could achieve 50%. At the end of experimental period, which was in 36-days, the average concentration of TN was just 0.58 mg/L. The effluent concentration of TN remained stable at the end of experimental period, especially after 24 days. The average concentration of NH_4_^+^-N could hardly be detected at the end of the experimental period and the average removal rate of NH_4_^+^-N could achieve 69%. The lower NH_4_^+^-N concentration caused by the small scale of drainage and release of wastewater from industrial production. The influent and effluent samples of COD were 29.628 and 10.714 mg/L, respectively. It indicated that MS-PFI could play obvious effect for the removal rate of COD were 61% in the experimental period.

**Fig 3.**
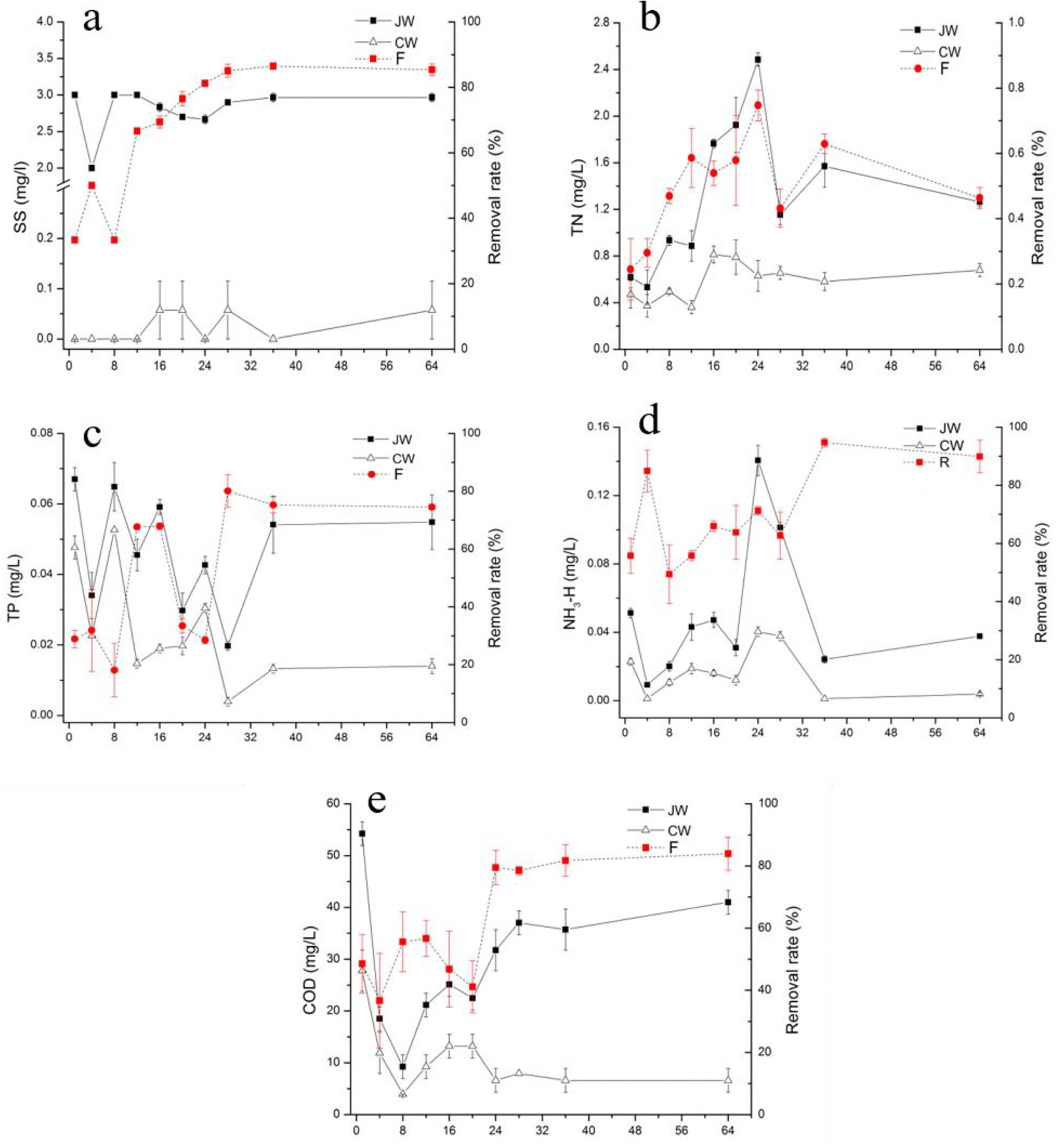
SS, TN, TP, NH_4_^+^-N, and COD were monitored in influent and effluent samples of MS-PFI

Ten samples were divided into three clusters, which was based on the removal rate of 7 water quality indexes (Fig.4). Blue cluster A grouped first-day, fourth-day and eighth-day samples based on the removal rates from 7 indexes, green cluster B aggregated twelfth-day, sixteenth-day and twentieth-day samples, red cluster gathered twenty-fourth-day, twenty-eighth-day, thirty-sixth-day and sixty-fourth-day samples. This indicated that the distribution of removal rates during different sampling time could reflect the water quality condition.

### Changes in the bacterial community

In this study, a total of 40 bacterial phyla across all samples were identified by using the OTU classifier (Fig. 5). Proteobacteria was one of the most abundant phyla in all samples, accounting for 40.7% and 46.1% of total influent and effluent effective bacterial sequences, respectively. Other dominant phyla were Actinobacteria (20.6 and 19.2%), Bacteroidetes (11.2 and 18.5%), Cyanobacteria (16.0 and 3.9%), Verrucomicrobia (4.1 and 1.4%), Saccharibacteria (0.6 and 4.3%), Firmicutes (2.7 and 1.3%), Chloroflexi (1.0 and 0.3%). Moreover, Chlamydiae, Parcubacteria, TM6, Acidobacteria, Armatimonadetes, Planctomycetes, Gemmatimonadetes, etc. are less than 5%. Most phyla existed in our study, was consistent with other pyrosequencing analyses of different types of constructed wetlands.

**Fig 4.**
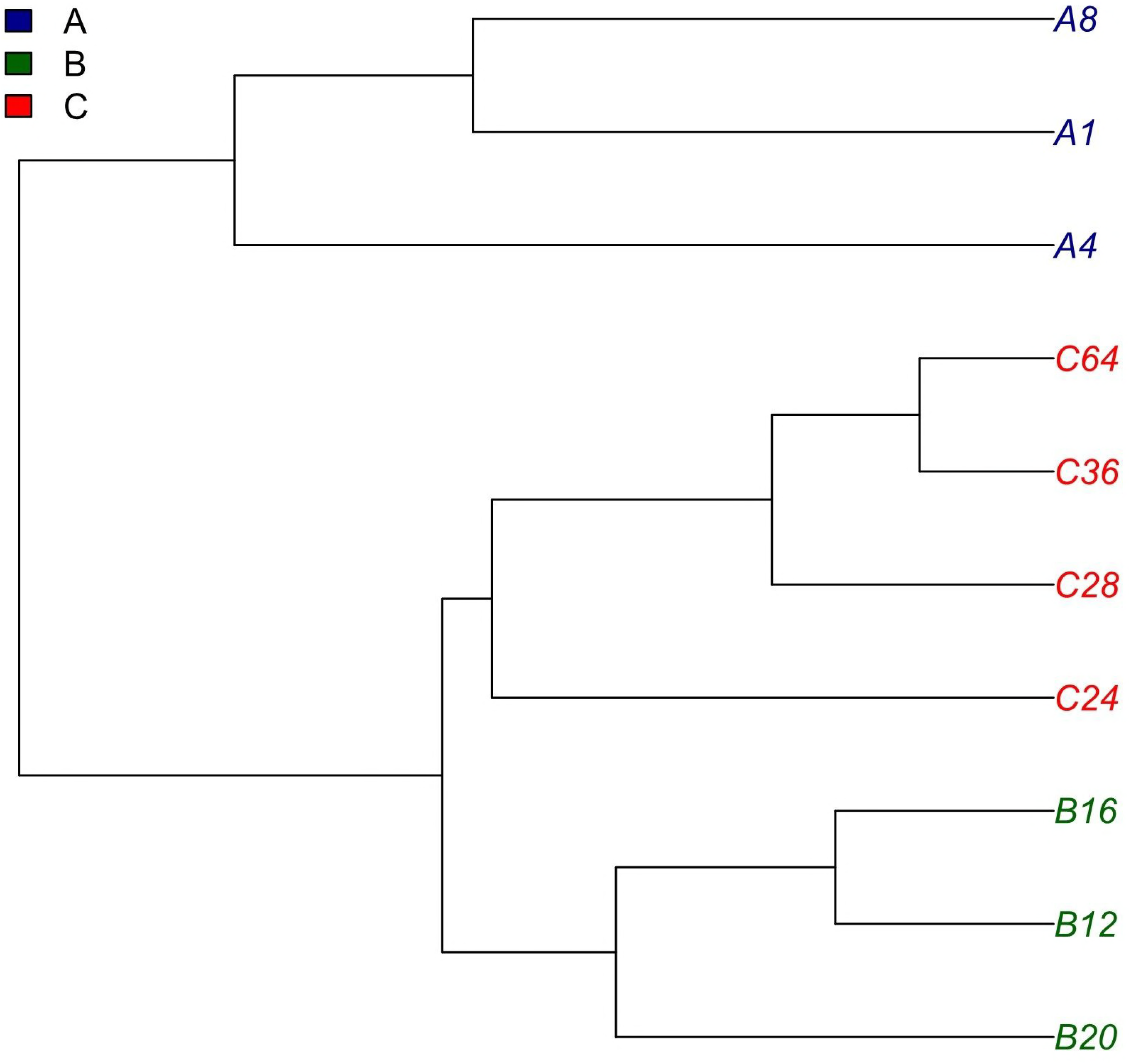
Hierarchical clustering analysis on removal rate of all samples based on 7 water quality indexes

**Fig 5.**
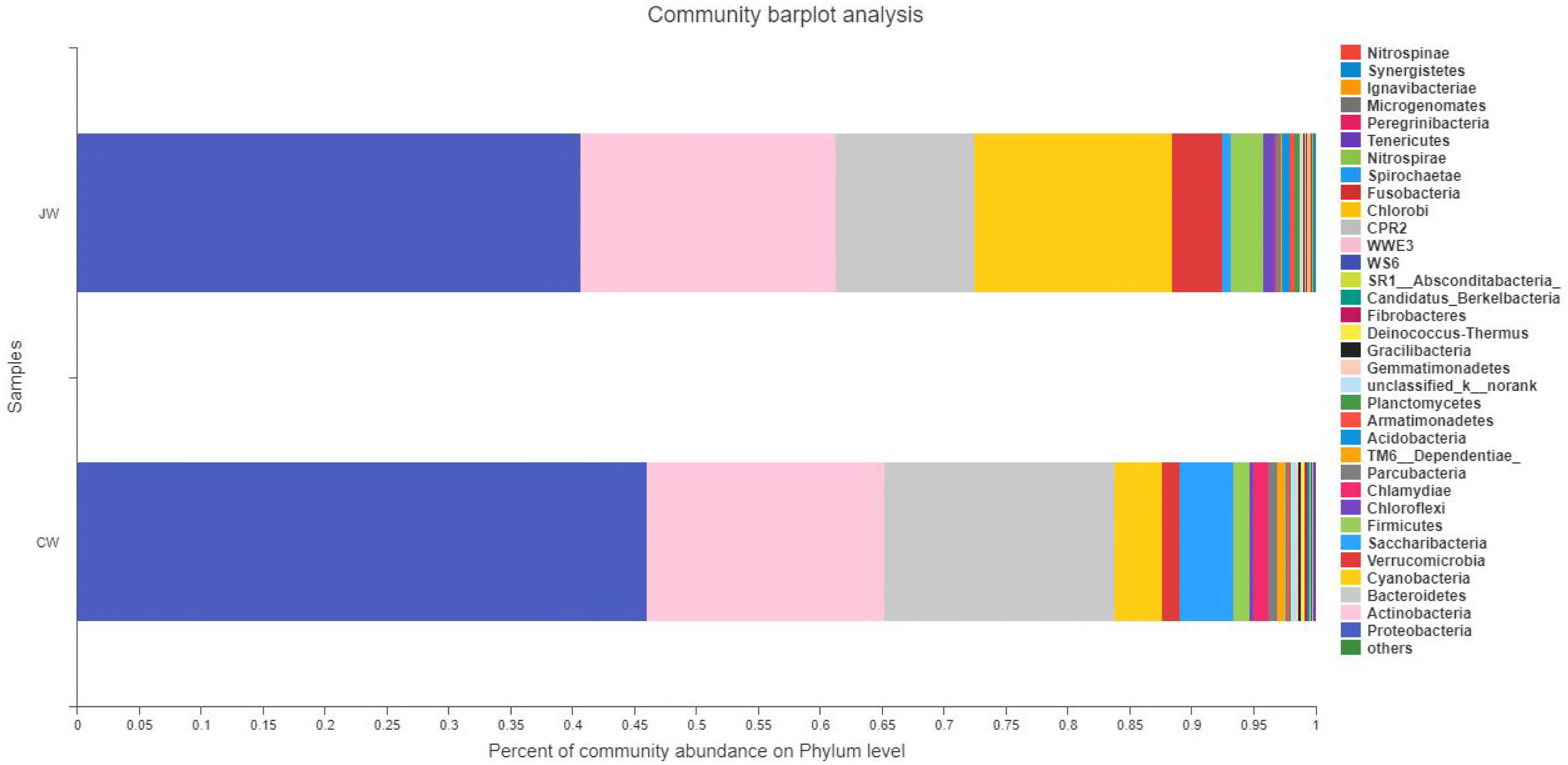
Bacterial community compositions at phylum level as revealed by pyrosequencing.

By characterizing the genera abundances in influent and effluent water of MS-PFI, metagenomic sequencing could unveil the variations of microbial richness and diversity in response to the environmental selective pressures (Fig. 6). For instance, 13, 12 and 7 % of SSU rRNA tags in JW were assigned to the hgcl_clade, Norank_c_Cyanobacteria and *Rhizorhapis*, while their relative abundances were decreased to 5, 3 and 0 %. A similar situation applied to abundances that were unclassified_f_Sporichthyaceae, norank_f_FamilyI, norank_f_Verrucomicrobiaceae and *Synechococcus*, Zymomonas, *Lactobacillus*, etc (Fig. 6). In contrast, the color intensity showed a distinct increase among *Flavobacterium* (10%), *Limnohabitans* (7%), *Alpinimonas* (4%), norank_p_Saccharibacteria (4%), *Erwinia* (3%). Moreover, microorganisms related to *Rhodococcus*, Candidatus_Planktoluna, *Pseudarcicella*, Candidatus_Rhodoluna, unclassified_f_ Comamonadaceae, *Sphingomonas*, *Novosphingobium*, *Rhizobium*, *Rhodobacter* enriched in the CW samples.

**Fig 6.**
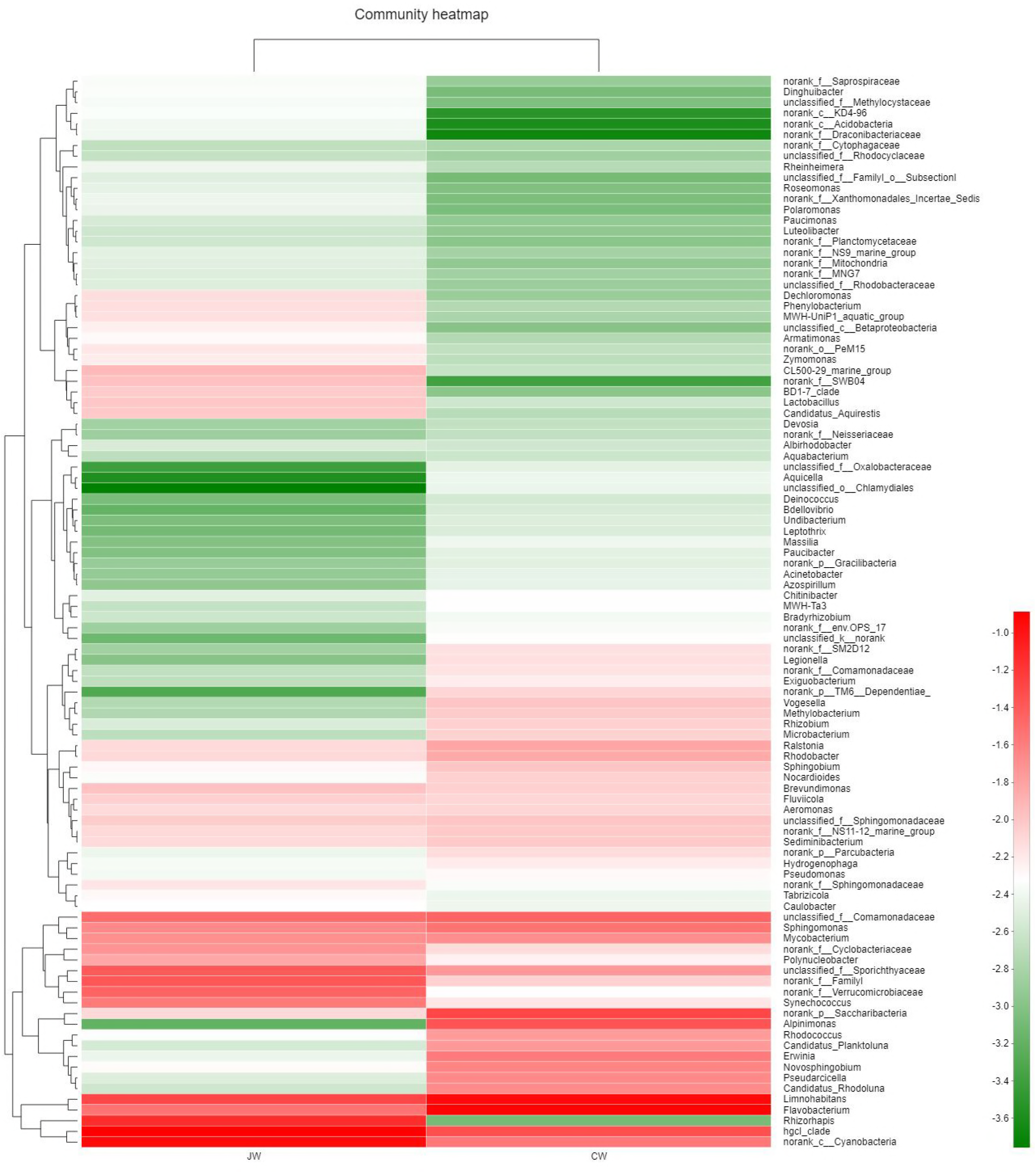
Heatmap of the top 100 bacteria genera for influent and effluent water samples were selected. The color intensity shows the abundance of genera as the color key indicates at the bottom right.

### Economic benefit evaluation of plant floating island

This study provides a common representative method of achieving buoyancy by polyethylene foam and acrylic sheets. Numerous commercial products available for achieving buoyancy or constructing the frame, especially a polystyrene foam, acrylic sheets, transparent hoses and foam mats, which is stable and cheap. And more importantly, this MS-PFI powered by solar energy, circled by pump and syphonage, without any exogenous energy investment and few equipment maintenances. The total cost of MS-PFI was calculated, which is shown in table 3. An amount of ¥ 1,843 was totally inputted to construct floating island, which included the cost of frame materials of floating bed, purchase of *Eichhornia* crassipes and expenses of labor.

**Table 3.** Economic benefit of plant floating island

| Items | Cost (¥) | Total cost (¥) |
| --- | --- | --- |
| *Eichhornia crassipes* | 100 |  |
| Solar energy pump | 260 |  |
| Accumulator | 238 |  |
| Acrylic sheets | 440 |  |
| Transparent hoses | 30 |  |
| PP, PVC, ABS, 502 glues | 85 | 1843 |
| Iron wires | 10 |  |
| EPE foam mats | 50 |  |
| Polystyrene foam | 100 |  |
| Ropes | 30 |  |
| Expenses of labor | 500 |  |

When maintenance was required, most jobs were replaced necrotic *Eichhornia* crassipes. In addition, during the fastest growing period, *Eichhornia* crassipes was occupied 95% of the water surface. Furthermore, plants could not only improve the water quality of draining channel but also provide a safe space for perching and living, for the best natural food natural for aquatic animals. Moreover, *Eichhornia* crassipes bloomed in May, it greatly increased the aesthetic value. The results indicated that enormous economic benefit could be achieved by this plant floating island. MS-PFI is a high-efficiency ecosystem, they can transform many inherent natural environmental energies, such as sun, wind, plant, and aquatilia, mounting fossil fuel energy, and chemicals will be replaced.

## Discussion

Our results and White’s study (4) indicated a significant difference between water inlet and outlet, effluent temperature was nearly 2°C lower the influent (Tab. 2). The decrease attributed to a higher percent coverage and the long retention time, which is a benefit for providing a healthy habitat by reducing the heat intake and reflecting most of heat. According to Rommens’ study (19), a remarkable difference existed in the present study, that is, pH value average increased by 0.11 after the MS-PFI treatment. Rezania et al. (20) indicated that a tiny growth in pH that ranges from 6.95 to 7.1 during an *Eichhornia* crassipes purification system. Some researchers considered that a decline of pH may drive by acid secretion from plant roots, and degradation of organic matter by aerobic organisms (21). However, others conceived that the ascent in pH attributed to blue algae growth by warming and a growth inhibition by NH_4_^+^ caused by saturation and depolarization of cell membranes.

The removal efficiency of the present study was higher than other field studies. For instance, the removal efficiency of TN, TP, NH_4_^+^-N, TDP, and chlorophyll-a were 36%, 35.7%, 44.3%, 38.1%, and 47.9%, respectively in Hu’s study (22). Moortel et al created an FTWs, showed lesser performance for TN, TP, NH_4_^+^-N and COD removal than present study (23). The removal efficiency of MS-PFI can be enhanced by adding biofilm carriers, immobilized microorganisms (15). For example, Song (24) added a biofilm carrier media, that performed greater effect for TN, TP and NH_4_^+^-N removal, 90.8%, 76.5% and 96.7% respectively.

Roots and polyethylene fillers play a vital role in reducing algae biomass and suspended particles, by physical filtration processes as well as chemical processes (25). Especially, the removal efficiency of SS was usually very high for filtration and sedimentation (9). Most P removals depended on SS settling, while N removal controlled by anaerobic ammonium oxidation, nitrification, denitrification and plant uptake (10). The high removal efficiency of NH_4_^+^-N and TN require a combination of HF (horizontal flow) and VF (vertical flow), namely hybrid MS-PFI, where each stage provides different redox conditions. The removal rate of COD related to the oxygen transfer rate (26), which based on aerenchymous tissues and leakage from roots. Fortunately, our experiments constructed a hybrid MS-PFI with a significantly effective.

The purification effect of MS-PFI made a difference during three periods and increased over time. Some researchers consider that *Eichhornia* crassipes absorbed nutrients on different stages and aerial tissues tend to have the highest nutrient concentrations. However, others (27) indicated that removal efficiency is enhanced regular harvesting, as the purification phase is followed by the decay phase. Based on this, our equipment would have a better removal efficiency by harvested once an annual as the animal food source.

Major phyla and the change trend of Proteobacteria were similar with Zhou’ study (12) in the *Eichhornia* crassipes treatments. Proteobacteria is an important microorganism involved in global carbon and nitrogen and sulfur cycle (28). Previous studies have revealed that although three major phyla (Protebacteria, Actinobacteria, Bacteroidetes) were detected in constructed wetlands samples, their relative abundances were different (29, 30). However, the results were different from the previous study reported by Yates et al. (31), who observed that Proteobacteria, Bacteroidetes, and Firmicutes. A wider range of phyla involved in wastewater treatment besides those mentioned above, for instance, Chloroflexi, Acidobacteria, Verrucomicrobia, Planctomycetes, Nitrospirae, Cyanobacteria, TM7 (also known as Saccharibacteria) or Gemmatimonadetes among others (28, 30, 32, 33).

*Zymomonas* and *Lactobacillus* are anaerobic, which confirmed that MS-PFI system may increase oxygen and inhibit anaerobic bacteria. A similar decrease related to *Dechloromonas* caused by the inhibition of oxygen. And it has been proposed as putative polyphosphate accumulating organisms (PAOs), accumulating polyphosphate and possibly contributing to denitrification (34). Reduction of the relative abundance of Norank_c_Cyanobacteria and *Synechococcus* occurred due to MS-PFI treatment. Zhou’ s study (12) indicated that *Eichhornia crassipes* not only introduced the root exudates but also had an inhibitory effect on cyanobacteria.

*Flavobacterium* and Comamonadaceae are ubiquitous microorganisms involving in contaminant degradation and nutrient removal in biological wastewater treatment systems (35). Homology search against NCBI *nr* database of the sequences assigned to this genus suggested that *Flavobacterium* were potentially associated with assimilatory and dissimilatory of nitrogen compounds which are preceded by NO_3_^-^/NO_2_^-^ transport into the cells. *Alpinimonas* isolated from river water and was likely to be important for dissolve organic matter removal (36). Norank_p_Saccharibacteria was a phylogenetically diverse group and played a role in the degradation of various organic compounds as well as sugar compounds under aerobic, nitrate reducing and anaerobic conditions (37).

*Rhodococcus* related to carbon and toxic compounds removal in Ferrera’ study (38). *Rhodobacter* is photoautotrophic without oxygen release, with H_2_ as the electron donor and CO_2_ as the electron acceptor and carbon source under anaerobic light or aerobic dark conditions (39). *Sphingomonas* has a great application potential for environmental protection and industrial production because of the high metabolic ability of aromatic compounds. *Novosphingobium* degraded aromatic compounds such as phenol, aniline, nitrobenzene, and phenanthrene (40). Moreover, microbial community indicated that less nitrifying bacteria (Nitrosomonasales and norank_c_Nitrospira, data were not shown) and more nitrogen-fixing bacteria (such as *Rhizobium*, *Bradyrhizobium*, and *Azospirillum*) after MS-PFI treatment provided it with a higher nitrogen recovery efficiency of *Eichhornia crassipes*. Consequently, elucidating the community changes and functional potentials of the microbial community is an important way to estimate the effect of water treatment.

Planted floating islands are considered a low-cost as well as an eco-friendly technology widely used to remediate water bodies in China (8,41), Europe (42), and America (4). The basic investment costs for MS-PFI include land, site investigation, system design, earthwork, liners, filtration or rooting media, plant, hydraulic control structures and fencing (26). Compared with Zhang’s study (7), we got a higher removal rate. And the capital, operations, and maintenance costs of our experimental system are about one-fifth of it. Moreover, our equipment rarely needs operation and maintenance costs, the costs are much lower than those for competing concrete and steel technologies (43).

## Conclusions

The plant floating island possesses the purification function and aesthetic values. Significantly, the set-up of MS-PFI vegetated with *Eichhornia* crassipes was efficient in the cleaning-up of waters polluted with SS, TN, TP, NH_4_^+^-N, and COD. The treatment efficiency and cost have directly limited the remediation of micro-pollution water, which influence surface water environmental quality. This system allowed the plants to grow and bloom, and then to operate a whole removal of pollutant within a month after the treatments, although pollutant concentrations were significantly lower in water inlet and outlet. Microbial community change caused by an increase of oxygen in MS-PFI system. It brings various bacteria involved in contaminant degradation and nutrient removal in biological wastewater treatment systems. These systems can be used in different environments and in combination with other traditional techniques to enhance the effectiveness and increase the potential of cleaning water bodies as landscape water, drainage channels, domestic, industrial and piscatorial wastewater, polluted groundwater, surface water, stormwater and irrigation water. Here, we provide another choice of MS-PFI with a concern for green energy and water quality maintenance.

## Materials and methods

### Plant floating island construction

The experimental units consisted of 3 experimental floating macrophyte wetlands that were constructed in three square troughs by top-down arrangement, with a length of 55 cm, 100 cm, 120 cm respectively and each trough 20 cm tall (Fig. 1). The bottom was fixed on the light polystyrene foam board to provide sufficient buoyancy (Conveying weight exceeded 50 kg). The trough was made of transparent acrylic sheets (Fig. 1-12), beautiful, low cost and easy-to-marketing application. *Eichhornia* crassipes (6.5 cm-tall) were rooted in buoyant foam mats (Fig. 1-3) floated on the surface of each trough. Each mat section had several (3 cm) pre-cut holes, which were spaced 5 cm on center. Each mat was designed to allow insertion of one plant seedling to avoid the phenomenon of dead plants caused by the high density (Fig. 1-11). The transparent latex tube (Fig. 1-5) used for these studies were 2 mm inner diameters, 4 mm outer diameter to ensure the siphon effect.

MS-PFI normal working was driven by solar energy (Fig. 1-2) and the solar panels installed on the top of MS-PFI (Fig. 1-14). Then sewage pumped to the trough top after flotages eliminated by the sieve (Fig. 1-6). Sewage was siphoned to the lower trough by siphon in trough wall. Sewage was circulated and purified in the trough through lifting, siphoning, and biodegradation, then discharged through the fountains (Fig. 1-7). It makes to fully improve water quality, also reduce the cost of aeration and less costly to install and maintain.

### Sampling sites and design

Experiments were conducted during the April 14 to June 16, 2017, each sample was set three repetitions. MS-PFI was placed near the drainage channel in Hefei University of Technology Xuancheng Campus (China, Anhui) and troughs were initially filled with sewage from it. Untreated sewage was collected from the water pump inlet and treated water was gleaned from the outlet of trough bottom. The samples were evaluated by sampling station and sampling time and were marked from JW1, JW4, JW8, JW12, JW16, JW20, JW24, JW28, JW36, JW64, and CW1, CW4, CW8, CW12, CW16, CW20, CW24, CW28, CW36, CW64 (JW means influent and CW implies effluent, Nos. 6 and 7 positions in Fig. 1) in sequence. Furthermore, sewage draining exit was collected from drainage channel as a reference sample. Water samples were collected using a 500ml water sampler (WB-PM, Beijing Pulite Instrument Co. Ltd., China).

### Physico-chemical variables analysis

The water samples were analyzed of water temperature (WT), pH, suspended solids (SS), total nitrogen(TN), total phosphorus (TP), ammonia nitrogen(NH_4_^+^-N) and chemical oxygen demand (COD_Cr_) both influent and effluent water samples. Water temperature was measured in situ by glass-stem thermometer (Sycif, Shanghai, China). All the other properties in water samples were determined according to the standard methods issued by the Chia State Environmental Protection Administration (44). Standard substances, guaranteed reagents, and other laboratory reagents were purchased from Sinopharm (Pharmaceutical group chemical reagent Co. Ltd., China.). All seven variables were determined completely on the day of sampling.

### DNA extraction, PCR amplification and High-throughput sequencing

DNA from the 18 samples (except JW64 and CW64) were amplified by PCR using primer set 338F (5’-ACTCCTACGGGAGGCAGCAG-3’) and 806R (5’-GGACTACHVGGGTWTCTAAT-3’) for the V3-V4 regions of the 16S rRNA (45). The reverse primer contains a 6-bp error-correcting barcode unique to each sample. The barcode was permuted for each sample and permitted the identification of individual samples within a mixture in a single Illumina MiSeq sequencing run. Approximately 200ml of water sample was used for DNA extraction with the E.Z.N.A.**^®^**. Water DNA kits D5525-01 (Omega, USA) according to the manufacturer’s instructions. Extracted DNA was and quantified using 1% agarose gel electrophoresis respectively, and then stored at −20 °C until use. PCR amplicons were further purified using a DNA purification kit (BioFlux, Japan), and the concentrations were determined by spectrometry using a QuantiFluor^™^ -ST (Promega, USA). Amplicons from different samples were then mixed to achieve equal mass concentrations in the final mixture, which were sent out to Majorbio Co., Ltd. In Shanghai for small-fragment library construction and pair-end sequencing using the Illumina MiSeq sequencing system (Illumina, USA)

### Bioinformatics analysis and data analysis

Pairs of reads from the original DNA fragments were merged using FLASH (46). Then the reads were filtered using QIME quality filters with the following criteria: (1) 250bp reads were truncated at any site receiving an average quality score<20 over a 50 bp by sliding window, and truncated reads shorter than 50 bp were discarded; (2) reads containing more than two nucleotide mismatches during primer matching or ambiguous characters were removed; (3) only sequences that overlap by more 10 bp were assembled according to their overlap sequence. Reads that could not be assembled were discarded.

Operational Taxonomic Units (OTUs) were clustered with a 97% similarity cutoff using UPARSE and chimeric sequences were identified removed using UCHIME (47). After the above filters were applied, the minimum number of selected sequences in the 18 samples was 685,656. The phylogenetic affiliation of each 16S rRNA gene sequence was analyzed by RDP Classifier (version 2.2 http://sourceforge.net/projects/rdp-classifier/) against the Silva (Release128 http://www.arb-silva.de) 16S rRNA database using a confidence threshold of 70%.

All data presented represent the average value for each sampling event ± the standard error of the mean. All plots were drawn by OriginPro 8.5.1 (OLC® 2011). Paired sample T-Test was performed with PASW Statistics 19.0.0 (IBM®, 2009) to analyze the difference of properties in the remediation and control areas. The significance level was set at P<0.05. Bar plot, hierarchical clustering analysis, and heatmap analysis were used by R. 3.3.1. Heatmap accounting for all consensus genera with top 50 relative abundance. Design drawing portrayed by CAD software.

## Acknowledgements

This work was supported by the Fundamental Research Funds for the Central Universities [grant number JZ2016HGTA0682], the doctoral industry-university-research of special funded projects from Hefei University of Technology Xuancheng Campus [grant number XC2016JZBZ08] and the Central Public-interest Scientific Institution Basal Research Fund, CAFS-ECSF (grant number 2018Z02-03). The authors would like to thank Di Enliang for portraying this design drawing.

